# The source of additive genetic variance of evolutionarily important traits

**DOI:** 10.1101/776898

**Authors:** Li Liu, Yayu Wang, Di Zhang, Xiaoshu Chen, Zhijian Su, Xionglei He

**Affiliations:** State Key Laboratory of Biocontrol, School of Life Sciences, Sun Yat-Sen University, Guangzhou, 510275, China; Zhongshan School of Medicine, Sun Yat-Sen University, Guangzhou, 510080, China; Department of Cell biology, Jinan University, Guangzhou, 510632, China

## Abstract

Fisher’s fundamental theorem of natural selection predicts no additive variance of fitness in a natural population. Consistently, observations in a variety of wild populations show virtually no narrow-sense heritability (*h*^2^) for traits important to fitness. However, counterexamples are occasionally reported, calling for a deeper understanding on the evolution of additive variance. In this study we propose adaptive divergence followed by population admixture as a source of the additive genetic variance of evolutionarily important traits. We experimentally tested the hypothesis by examining a panel of ~1,000 yeast segregants produced by a hybrid of two yeast strains that experienced adaptive divergence. We measured over 400 yeast cell morphological traits and found a strong positive correlation between *h*^2^ and evolutionary importance. Because adaptive divergence followed by population admixture could happen constantly, particularly in some species such as humans, the finding reconciles the observation of abundant additive variances in evolutionarily important traits with Fisher’s fundamental theorem of natural selection. It also suggests natural selection may effectively promote rather than suppress additive genetic variance in species with wide geographic distribution and strong migratory capacity.

## Introduction

A basic issue of evolutionary genetics is to understand the relationship of natural selection and fitness (Orr 2009; Hendry, et al. 2018). The Fisher’s fundamental theorem of natural selection predicts little additive genetic variance (or narrow-sense heritability, *h*^2^) for fitness, because natural selection will fix alleles with the highest fitness quickly (Mousseau and Roff 1987; Merila and Sheldon 1999a; Crow 2002). An extended prediction of the theorem is that traits tightly-coupled with fitness (i.e., evolutionarily important traits) should have smaller *h*^2^ than those less-coupled with fitness (Kruuk, et al. 2000), because the response to natural selection on fitness will shape the evolution of the related traits (Orr 2009). The negative correlation between *h*^2^ and trait importance has been found in a variety of studies on different species and/or populations (Merila and Sheldon 1999b; Kruuk, et al. 2000; Merila and Sheldon 2000; Stirling, et al. 2002; Teplitsky, et al. 2009; Wheelwright, et al. 2014; Sztepanacz, et al. 2017). For example, for the wild female red deer (*Cervus elaphus*) the *h*^2^ of several life history traits, including the total number of offsprings, the adult breeding success and the longevity, were all zero (Kruuk, et al. 2000). Meanwhile, the morphologic traits such as leg length and jaw length, which are believed to be less related to fitness, were found to have much higher *h*^2^ than the life history traits. The pattern is also true for collared flycatcher (*Ficedula albicollis*), Savannah sparrows (*Passerculus sandwichensis*), red-billed gull (*Larus novaehollandiae*), and so on (Merila and Sheldon 2000; Stirling, et al. 2002; Teplitsky, et al. 2009; Wheelwright, et al. 2014).

However, there are also reports of abudant additive variances for important traits (Pettay, et al. 2005; Teplitsky, et al. 2009; Kosova, et al. 2010; Milot, et al. 2011; Zhang 2012). In particular, there is even a positive correlation between *h*^2^ and trait importance. For example, in a bighorn sheep population from Ram Mountain the lowest *h*^2^ was for body mass at primiparity (0.02), while the *h*^2^ of lifetime fecundity was as high as 0.66 (Reale and Festa-Bianchet 2000). A variety of explanations to the observations have been proposed. In addition to considering the definition of fitness, the variance components of *h*^2^ and balancing selection, a predominant view is that fluctuating environments combined with mutations could help maintain high additive variance of fiteness (Burger and Gimelfarb 2002; Crow 2008; Zhang 2012). These explanations are all theoretical, lacking empirical evidence. More importantly, they do not predict a positive correlation between *h*^2^ and trait importance.

We reason that here an ecological factor in evolution, namely, migration, may play an essential role. For a given species there are often plenty of divergences among populations (Pizzo, et al. 2008; Sved, et al. 2008; Roy, et al. 2014). When the divergences are coupled with local adaptation (i.e., adaptive divergence), which happens quite often in nature (Pulido 2007; Liedvogel, et al. 2011), alleles with beneficial additive effects on important traits would be preferentially fixed in a population. Since different genes would be selected for in different populations, subsequent population admixture by migration would lead to a new population with abundant additive genetic variances for important traits. In this study we designed an experimental test for this theoretically compelling hypothesis.

## Results

We started from a panel consisting of ~1,000 prototrophic haploid yeast segregants produced from a cross of two *Saccharomyces cerevisiae* strains (BY parent and RM parent) that were subject to adaptive divergence (Ho, et al. 2017). The two parental strains are diverged by ~0.5% at the genomic sequence level and the segregants were all genotyped in a previous study (Bloom, et al. 2013). We first verified the segregant panel and removed the segregants that appeared to be discordant with the reported genotypes (Fig. 1A, Methods).

**Figure 1.**
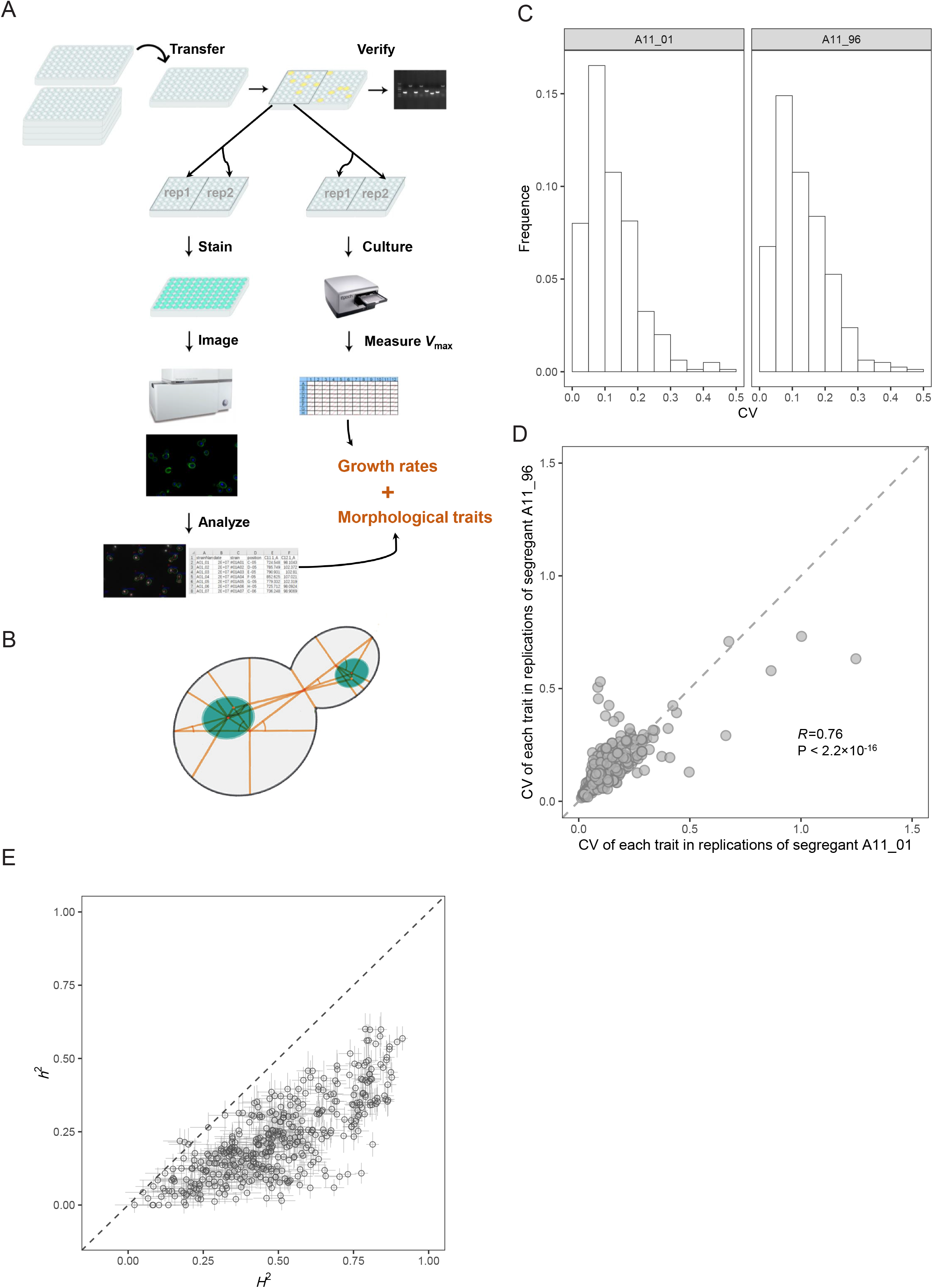
Measuring 405 cell morphological traits and their heritability in the yeast segregant panel. **(A):** The experimental process in this study. Both morphological traits and growth rate of each segregant were measured with replications. **(B):** The schematic diagram of calculating morphological traits. **(C):** The distributions of CV of each morphological trait calculated by replications in A11_01 (left) and A11_96 (right), respectively. **(D):** The relationship of CV of each morphological trait between A11_01 and A11_96 (Pearson’s *R* = 0.76, *P* < 2.2×10^−16^, *N* = 405). **(E):** The broad-sense heritability (*H*^2^) and narrow-sense heritability (*h*^2^) for each trait. Error bars represent SE, and the dashed line represents *h*^2^ = *H*^2^ (Pearson’s *R* = 0.77, *P* < 2.2×10^−16^, *N* = 405).

We measured 405 cell morphological traits for each segregant with two technical replications by following a previous protocol with some modifications (Methods) (Ohya, et al. 2005). These traits are related to the characters of mother cell and/or bud in different stages, such as area, distance, localization, angle, ratio and so on (Fig. 1B). After excluding measurements with insufficient cell number for calculating traits (Methods), we obtained the morphological trait information for 734 segregants, 73.3% (538/734) of which had data of at least two replications (Table S1). Approximately 99.5% of the trait values were derived from over 100 cells of a given segregant (Fig. S1). Segregants A11_01 and A11_96 were measured in every experiment as a technical control for potential operating bias in culturing, staining, and imaging. The coefficient of variance (CV) of each trait is rather small, of which 80% are less than 0.2 in both segregants (347/405 in A11_01, 326/405 in A11_96, Fig. 1C). In addition, CVs of the 405 traits are highly correlated between A11_01 and A11_96 (Pearson’s *R* = 0.76, *P* < 2.2×10^−16^; Fig. 1D). These data together suggest no strong batch effects in the trait measurements. We also checked pairwise rank correlation of the 405 traits between 28 technical replications of the segregant A11_96, or 26 replications of the segregant A11_01, or two replications of 536 segregants, respectively. We observed invariantly strong correlations, suggesting the reliability of trait measurements (Fig. S2). The large number of high-quality quantitative traits of the same property (i.e., morphology) measured under the same experimental setting provide a unique opportunity to study the evolution of additive genetic variance.

Quantile normalization of the raw trait values was performed to ensure the different traits comparable (Methods). The broad-sense heritability *H*^2^ and narrow-sense heritability *h*^2^ were estimated for each of the 405 traits according to an previous study (Bloom, et al. 2013) (Table S2). Notably, there are neither dominance effects since the segregants are haploid nor gene-environment interactions since they all grow in the same environment. Thus, in this study *H*^2^ includes additive effects and between-gene epistatic effects and *h*^2^ includes only additive effects. The *H*^2^ of the 405 traits ranges from 0.021 to 0.913, with a median of 0.478; the *h*^2^ ranges from 0.000 to 0.619, with a median of 0.240 (Fig. 1E). As expected, there is a strong positive correlation between *H*^2^ and *h*^2^ (Pearson’s *R* = 0.77, *P* < 2.2×10^−16^).

To assess the evolutionary importance of the morphological traits we computed their relatedness to <u>g</u>rowth rate (RTGR). We measured the growth rate of each segregant under the same condition as for trait measurement (Fig. 1A, Methods, Table S3) (Orr 2009; Chen, et al. 2017; Hendry, et al. 2018). For each of the 405 traits we computed the Pearson’s correlation coefficient (Pearson’s *R*) between trait value and cell growth rate among the 734 segregants. Following a previous study (Chen, et al. 2017), the absolute value of Pearson’s R was then used as the RTGR of a morphological trait; traits with larger RTGR are regarded as evolutionarily more important. The value of RTGR varies from 0 to 0.308, with a median of 0.065, highlighting a wide range of evolutionary importance of the 405 morphological traits (Fig. 2A).

**Figure 2.**
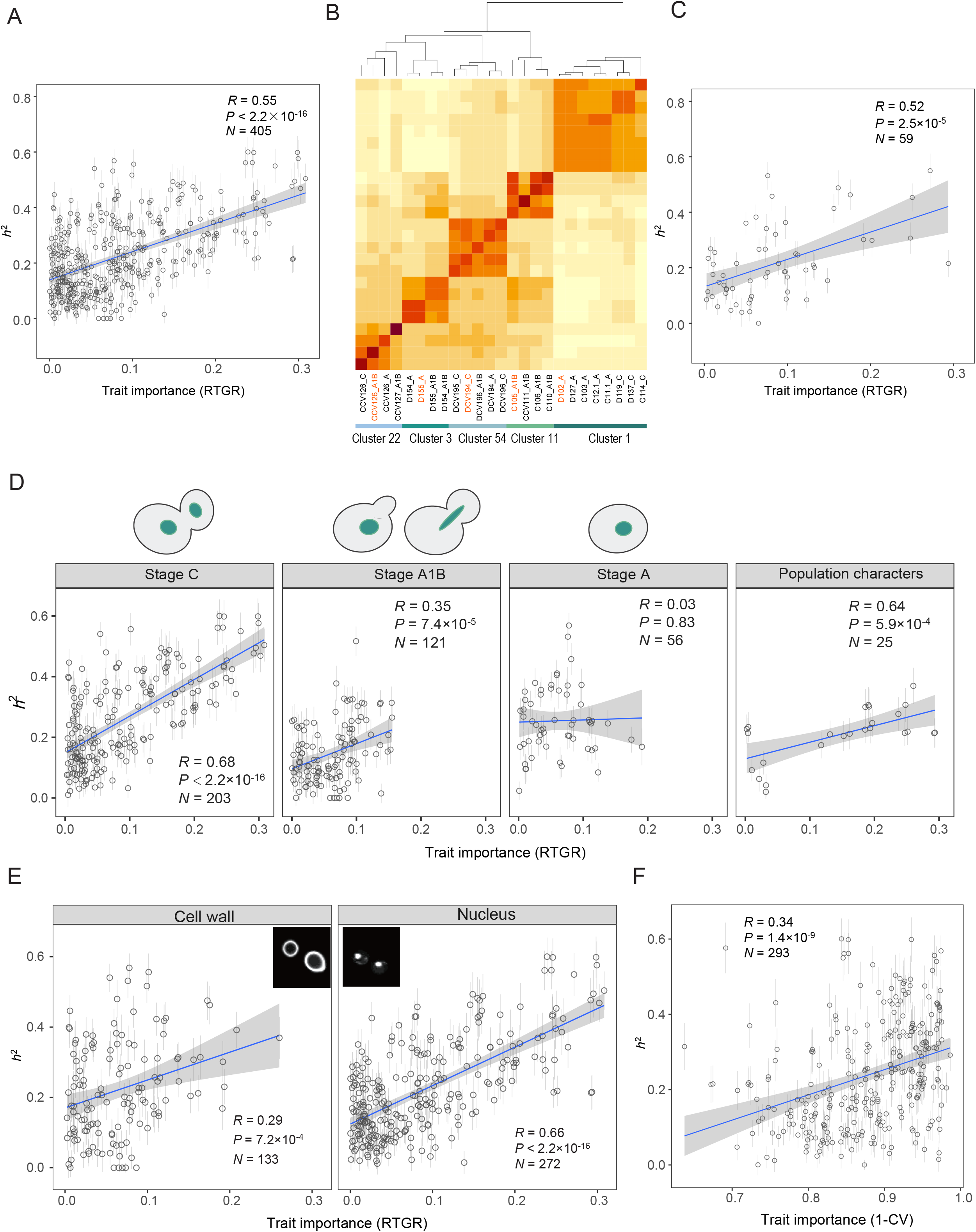
A positive correlation between trait importance and *h*^2^ among the 405 morphological traits. **(A):** A strong positive correlation between *h*^2^ and trait importance estimated by RTGR across 405 morphological traits (Pearson’s *R* = 0.55, *P* < 2.2×10^−16^, *N* = 405). Error bars represent SE. The blue line is the linear regression line and the gray zone shows the 95% confidence interval. **(B):** A heatmap of similarity matrix for traits in Cluster 1, 3, 11, 22, 54 derived by apcluster, as an example of obtaining exemplary traits. The top lines show the clustering results, and the corresponding traits are marked below. The exemplary trait in each cluster is colored as orange. **(C):** The strong positive correlation between *h*^2^ and trait importance estimated by RTGR across exemplary traits which are less correlated to each other (Pearson’s *R* = 0.52, *P* = 2.5×10^−5^, *N*= 59). **(D):** The correlation between *h*^2^ and RTGR for traits characterized cell morphology at different cell cycle stages. The top cartoons show the states of mother cell and bud of each category. **(E):** The correlation between *h*^2^ and RTGR for traits characterized different cell components. Traits derived from cell wall were stained by FITC-ConA, and traits related to nucleus were stained by hoechst. **(F):** A positive correlation between *h*^2^ and trait importance estimated by 1-CV across 298 morphological traits (Pearson’s *R* = −0.37, *P* = 5.3×10^−11^, *N* = 293).

According to our hypothesis, the admixture of two populations with adaptive divergence would result in a new population with more additive variances in evolutionarily more important traits. The availability of both *h*^2^ and evolutionary importance for the large number of traits enables a direct test for the hypothesis. Consistent with the hypothesis, we found a strong positive correlation between *h*^2^ and trait importance estimated by RTGR among the 405 yeast traits (Pearson’s *R* = 0.55, *P* < 2.2×10^−16^; Fig. 2A). Because many traits are correlated with each other, we conducted affinity propagation clustering and obtained 59 trait clusters each with an exemplary trait (Fig. 2B and Fig. S3, Methods). The number of traits represented by an exemplary trait ranges from 2 to 16, with a median of 6, and there are only weak correlations among the 59 exemplary traits (Fig. S3). The strong positive correlation between *h*^2^ and RTGR remained when only the exemplary traits were considered (Pearson’s *R* = 0.52, *P* = 2.5×10^−5^, Fig. 2C).

The 405 traits represent cell morphology at different cell cycle stages. We divided these traits into four categories according to the states of bud and nucleus (Methods). Traits of stage A1B and stage A tend to have small RTGR, suggesting less selective constraints on the morphology of the two stages. Importantly, the positive correlation between *h*^2^ and RTGR remained with the exception for traits of stage A (Fig. 2D). In addition, since the 405 traits represent features of cell wall and nucleus that are stained by two different dyes FITC and Hoechst, respectively (Methods), we examined the 133 cell wall-related traits and 272 nucleus-related traits separately. The positive correlation between *h*^2^ and RTGR holds in both categories (Fig. 2E).

A previous study suggests the coefficient of variance (CV) in trait measurement could serve as a proxy of trait importance, with smaller CV for more important traits (Ho and Zhang 2014). We thus calculated the measuring CV of each morphological trait using the 26 replications in segregant A11_01, and the 28 replications in segregant A11_96, respectively. To be conservative we considered only 293 traits that have consistent CV between A11_01 and A11_96, and used the average to represent trait importance (Methods; Fig. S4). We observed a positive correlation between *h*^2^ and 1-CV (Pearson’s *R* = 0.34, *P* = 1.4×10^−9^; Fig. 2F), a result supporting our hypothesis. The pattern holds by considering exemplary traits or by separating the traits into different categories (Fig. S4).

We then mapped quantitative trait loci (QTL) for each of the traits. A total of 2,505 QTLs were detected for 393 traits (Table S4), and the number of QTLs ranged from 1 to 19, with a median of 6 for each trait (Fig. S5). There are 12 traits with no detectable QTLs, which conforms to their extremely low *h*^2^ (median *h*^2^ ~ 0.016). In nearly all cases the trait variance explained by detected QTLs is close to *h*^2^ (Pearson’s *R* = 0.96, *P* < 2.2×10^−16^, Fig. 3A), suggesting limited missing heritability. This is consistent with a previous observation in the yeast segregant panel (Bloom, et al. 2015). Most of the QTLs (~90.6%) each explains less than 5% trait variance (Fig. S5).

**Figure 3.**
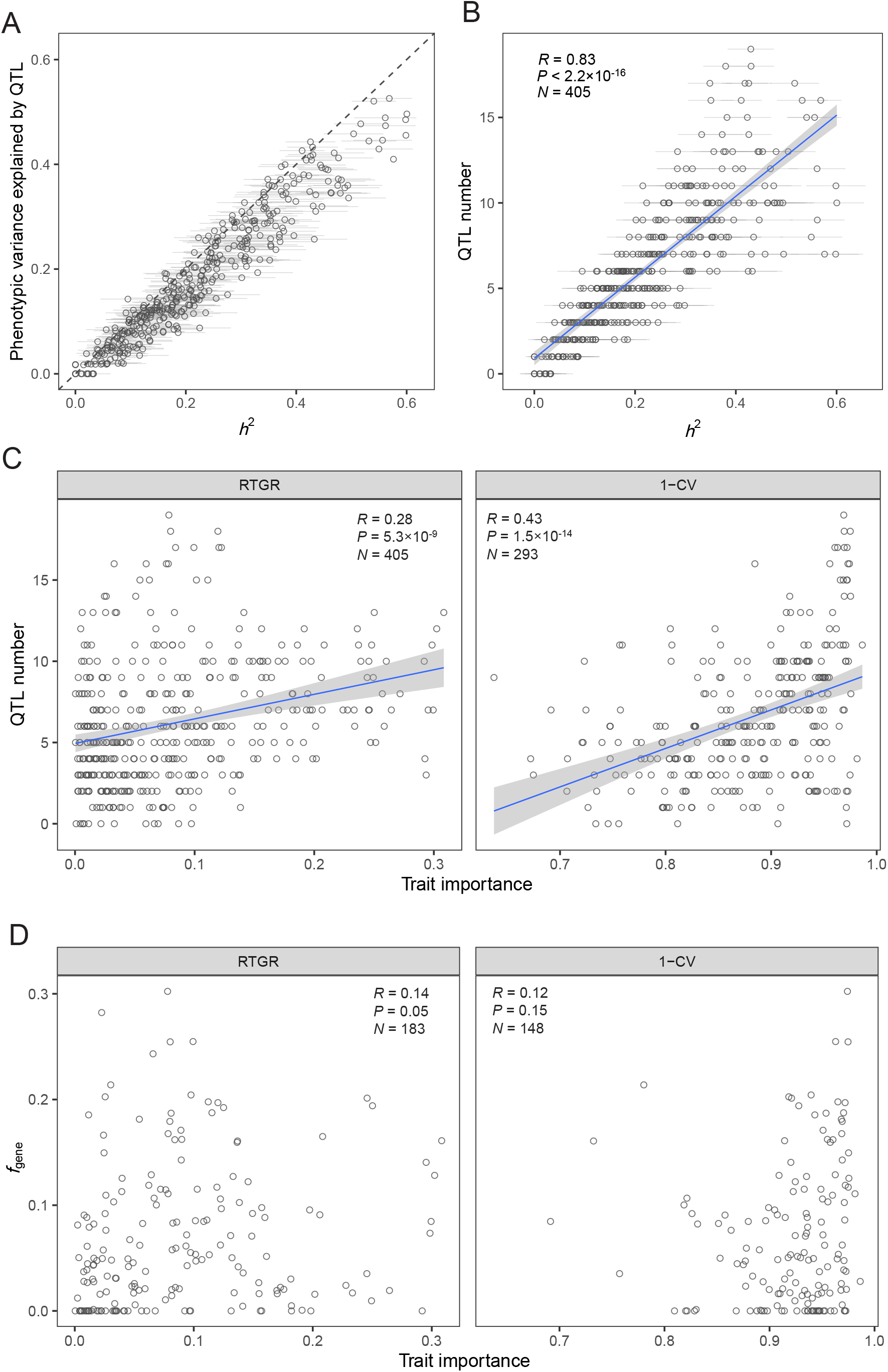
Analysis of QTLs of the 405 morphological traits. **(A):** The narrow-sense heritability (*h*^2^) of each trait is plotted against the phenotypic variance explained by additive QTLs (Pearson’s *R* = 0.96, *P* < 2.2×10^−16^, *N* = 405). Error bars represent SE, and the dashed line represents variance = *h*^2^. **(B):** A strong positive correlation between QTL number and *h*^2^ (Pearson’s *R* = 0.83, *P* < 2.2×10^−16^, *N* = 405). The gray zone shows the 95% confidence interval of the regression line (blue). **(C):** Positive correlations between QTL number and trait importance estimated by RTGR (left) and 1-CV (right). **(D):** No apparent correlation between the fraction of genes that affect a trait (*f*_gene_) and trait importance was observed. Only a proportion of traits have estimated *f*_gene_ values.

We found the *h*^2^ of a trait is highly correlated with the number of QTLs (Pearson’s *R* = 0.83, *P* < 2.2×10^−16^; Fig. 3B). Consistently, there are in general more QTLs detected for more important traits (Fig. 3C and Fig. S6). This result means there are more diverged loci for important traits after the split of the two parental yeasts of the segregant population examined here. There are two possible explanations: First, there are more genes and thus more mutations that affect important traits; second, there are higher fixation rates for mutations that affect important traits. To answer the question, we examined the cell morphology data generated for a large set of yeast single gene deletion mutants. For each of the traits we obtained the fraction of genes that affect a trait (*f*_gene_) by following a previous study (Ho and Zhang 2014). We failed to observe a larger *f*_gene_ for more important traits (Fig. 3D), suggesting the second explanation is plausible although the number of whole genes affecting a trait does not necessarily tell the number of natural variants affecting the trait. A higher fixation rate of mutations affecting more important traits indicates positive selection underlies the genetic divergences of the parental yeasts, echoing the adaptive phenotypic evolution of the yeasts previously proposed based on the faster evolution of more important traits within and between species (Ho, et al. 2017).

## Discussion

Fisher’s fundamental theorem of natural selection provides a general framework for thinking of the evolution of additive genetic variance. Previous empirical studies on this issue are all based on wild populations and the resulting patterns are discordant, which are often ascribed to confounding ecological factors. This study is, to the best of our knowledge, the first controlled experiment for examining the relationship between additive variance and evolutionary importance in a large set of quantitative traits. The advantage of the controlled experiment is the ecological variables common to individuals of a wild population, such as nutrition, parasite, predator, and so on, are all fixed. However, there is a caveat in the design. Specifically, since the proposed adaptive divergences of the two parental yeasts must occur in specific natural environments, the trait importance obtained in the lab condition may not necessarily represents that of the environments. Nevertheless, this problem would more likely reduce the signal of the positive correlation between *h*^2^ and trait importance.

Our finding has implications for understanding the origin and maintenance of additive genetic variances. This is particularly true for humans who have both wide geographic distribution and strong migratory capacity, the former predicting frequent local adaptions (i.e., adaptive divergences) and the latter enabling constant population admixtures (Hellenthal, et al. 2014; Fu, et al. 2016). During adaptive divergence the additive genetic variances would be preferentially fixed by positive selection, which is strongest for the most important traits. Because different genetic loci would be selected for in different populations, subsequent population admixture would lead to a new population with abundant additive genetic variances. The process of adaptive divergence followed by population admixture could be repeated again and again during the human evolution (Fig. 4), resulting in the current population structure in which positive selection may have effectively promoted rather than suppressed additive genetic variances. This proposal might be of great value for understanding the genetic architecture of human complex traits.

**Figure 4.**
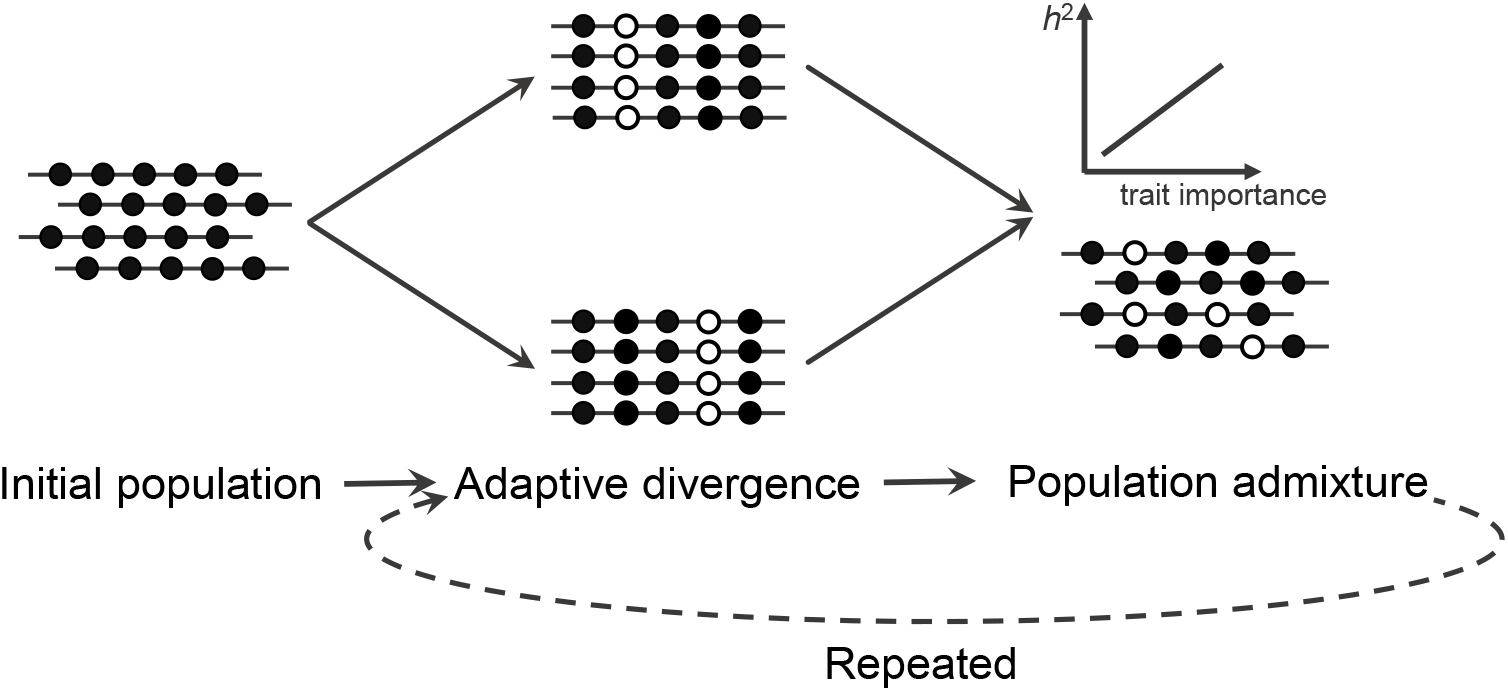
A model of adaptive divergence followed by population admixture underlies the origin and maintenance of additive genetic variances. The genotype of an initial population is assumed to be uniform. During adaptive divergence, different genetic loci would be selected in different populations, subsequent population admixture would lead to a new population with abundant additive genetic variances, and more important traits would have more additive variances.

## Methods

### Verify segregant panel

The segregant panel was kindly provided by Dr. L. Kruglyak. There were total 1,056 segregants in eleven 96-well plates. To verify the genotypes, twelve segregants in each plate were randomly picked up and four loci (MATa, MATα, hphMX4, natMX4) were amplified by polymerase chain reaction (PCR) for these segregants. By comparison the results with the genotypes provided by Dr. L. Kruglyak, we found that some percentage of segregants in Plate 8 and 9 were mismatched, and there was no pattern to rescue the segregants in a row or a line, which may be the result of contaminations. We then focused the segregants in the left nine plates with right genotypes in the next experiments.

### Measure cell morphological traits

The morphological traits of each segregant were measured following Ohya’s protocol with some modifications (Ohya, et al. 2005). Briefly, segregants were grown in YPD medium (yeast extract/peptone/dextrose medium) to saturation phase at 25°C for two or three days, and then transferred to new cultures to exponential phase at 25°C for three or four hours. Each segregant had two replications. Cells were fixed with 3.7% formaldehyde solution. Cell walls were stained with FITC-ConA (fluorescein isothiocyanate-conjugated, concanavalin A, Sigma-Aldrich C7642). Cell nucleus were stained by hoechst-mix (Thermo Fisher, Hoechst 33342 Solution) instead of DAPI to enhance the specificity. We omitted the process of actin staining because the dye of actin (Rhodamine phalloidin) was not stable and couldn’t support to image for a long time in the high-throughput automated image-processing. The stained cells were plated on microplates (Greiner 781091) with ~5.0×10^4^ cells per well and taken images by IN Cell Analyzer 2200 (GE Healthcare) with 100× objective lens. There were two technical replications for each segregant, and segregants A11_01 and A11_96 were cultured, stained, and imaged in every experiment as a technical control.

CalMorph software was used to analyze images to quantify yeast morphology, and 405 quantitative traits were derived (Ohya, et al. 2005). Segregants whose cell number for calculating traits less than 80 in both two replications were excluded. Values of all traits were listed in Table S1. There were 734 segregants each with 405 morphological traits derived, in which 73.3% (538/734) had at least two replications. Quantile normalization was performed to the raw values of traits by R package preprocessCore for further calculations (Bolstad 2019).

Traits derived from cell wall or nucleus can be distinguished by the initial letter of traits, which “C” is related to cell wall, and “D” is related to nucleus. Traits in different stages can be distinguished by the letters after the connector line. “A” represents traits calculated by cells with one nucleus and without a bud, “A1B” is traits calculated by cells with one nucleus in the mother cell with a bud or the nucleus is dividing at the neck, and “C” is traits derived by cells with one nucleus each in the mother cell and bud. The 405 traits are not independent, and 59 exemplary traits were derived by R package ‘apcluster’ (negDistMat, r = 2) using the mean normalized values of 734 segregants (Frey and Dueck 2007).

### Measure cell growth rate

Strains were grown in YPD medium to saturation phase at 25°C for two or three days, then diluted 1:100 to 100ul fresh YPD medium at 96-well plate. Two replications of each segregant were placed in the same 96-well plate. The 96-well plates were put on Epoch2 Microplate Spectrophotometer (BioTek) and incubated at 25°C with shaking. The absorbances at 600 nm of each well were determined per hour. The measurements lasted 24 hours and all strains reached saturation phase. The Vmax of growth rate, i.e. the maximum slop of growth curve of each well, was used to estimate the fitness of each strain. To control the positional bias, the original values of growth rate in each plate were fit by a robust locally weighted regression by R package ‘locfit’ according to Bloom et al’s study (Bloom, et al. 2013). The average normalized values of growth rate were taken as the fitness of each segregant, and listed in Table S3.

### Calculate heritability

Because the segregant panel was produced by Bloom et al, broad-sense heritability (*H*^2^), narrow-sense heritability (*h*^2^), additive QTL, and the variance explained by QTL of each morphological trait were calculated by methods consisted with Bloom et al’s study (Bloom, et al. 2013). Briefly, *H*^2^ was calculated by normalized values of traits of segregants with two replications. *H*^2^ was estimated as 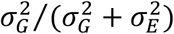, where 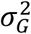 was the genetic variance and 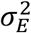 was the error variance. It was performed by the ‘lmer’ function in lme4 R package (Bates, et al. 2015). When compared to *H*^2^, *h*^2^ was calculated by the average normalized values of traits of segregants with two replications. And segregants with only one replication were also included in other situations. Narrow-sense heritability was estimated as 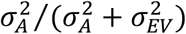, where 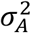 was the additive genetic variance and 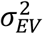 was the error variance. R package rrBLUP was used to calculate *h*^2^ (Endelman 2011). Standard errors of *H*^2^ and *h*^2^ were calculated by delete-one jackknife both.

Additive QTL of each trait was detected using the step-wise forward-search approach developed by Bloom et al (Bloom, et al. 2013). Lod scores for each genotypic marker and each trait were calculated as 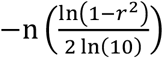, where *r* is the Pearson correlation coefficient between the genotypes and trait values. Significant genetic markers were detected from four rounds using different lod thresholds corresponding to a 5% FDR, which were 2.68, 2.92, 3.72 and 4.9, respectively. A multiple regression linear model was estimated by taken each QTL as independent variables of each trait, and the total phenotypic variance explained by additive QTL was the square of the multiple regression coefficient. The results were listed in Table S2.

### Calculate trait relatedness to growth rate

For each trait, the average normalized values of replications of each segregant were calculated. For segregants with only one replication, the trait values were the normalized values from the only measurement. The absolute Pearson’s *R* between the trait values and the growth rates in YPD medium across 734 segregants was used as a proxy of relatedness to growth rate for each trait. The results were listed in Table S2.

### Calculate trait CV among replications

Coefficient of variations for each trait were calculated using raw data of replications of A11_01 and A11_96, respectively. Traits with large CV may indicate their indeterminacy, so we excluded 7 traits with CV larger than 0.4 when using CV as an index of trait importance. To evaluate the repeatability of two groups, we used a distance index between two groups of CV as 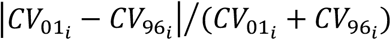, where 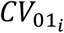 and 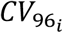 were the value of CV for trait *i* in A11_01 and A11_96, respectively. A large CV distance means the environmental robustness of the trait would be different for different segregants, so we excluded 105 traits with CV distance index larger than 0.2 to obtain a set of traits with consistent measurements. There were 293 traits left. The results were listed in Table S2.

## Acknowledgment

The yeast segregant panel is gift from Dr. Kruglyak at UCLA. This work is supported by research grants of NSFC (grant #31630042 and #91731302 to X. H.).

## Legends for supplementary figures

**Figure S1.**
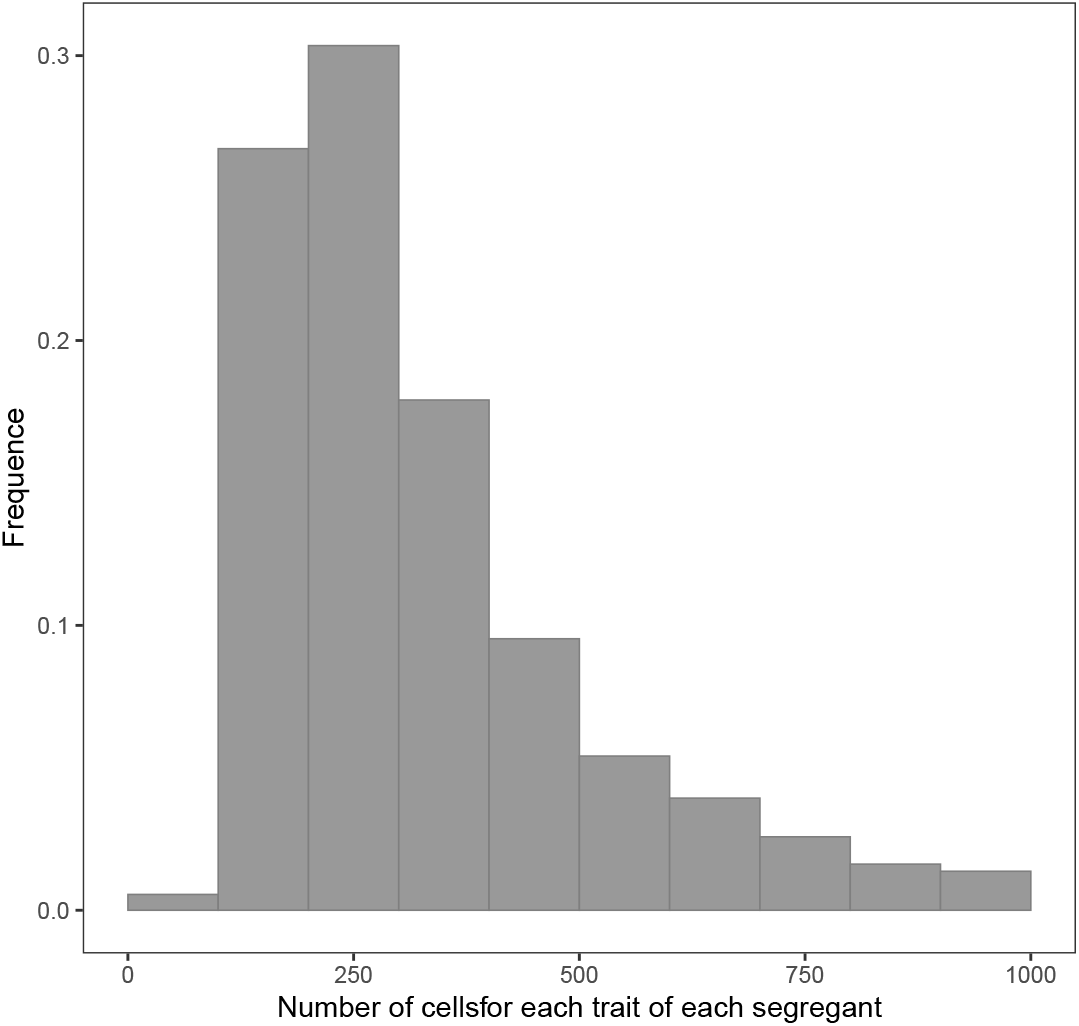
The distribution of cell numbers used to calculate each trait in each segregant.

**Figure S2.**
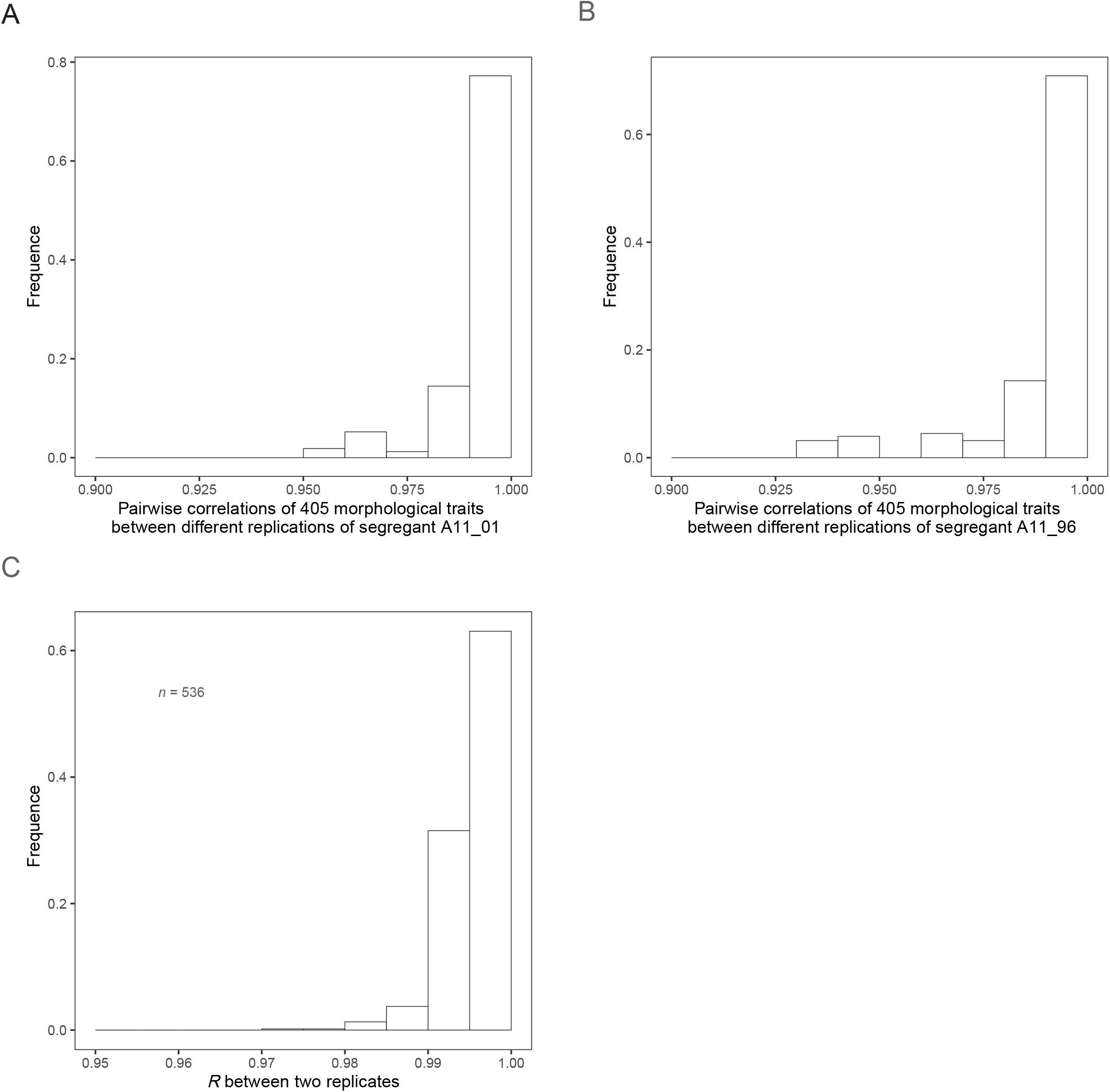
The distributions of pairwise correlations of 405 morphological traits between different replications of segregant A11_01 (A), A11_96 (B), and segregants with two replications (C). For A11_01, the spearman correlation of trait values between two replications was calculated, and this process was traversed for all replications. The same calculations were performed for A11_96. For the left 536 segregants each with two replications, the spearman correlation between two replications for each segregant was calculated.

**Figure S3.**
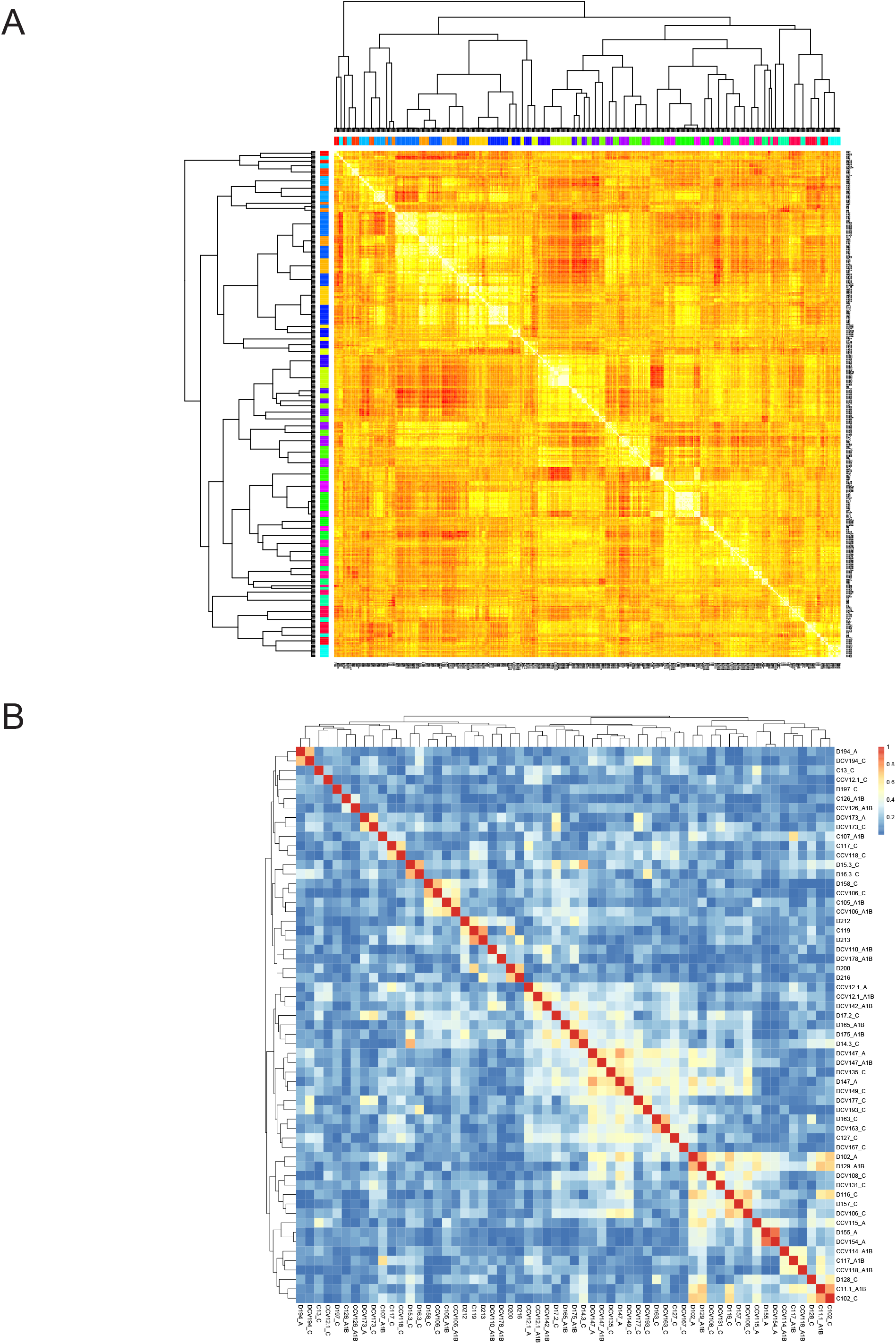
Fifty-nine exemplary traits were derived. (A): the similarity matrix of 405 traits derived by apcluster. (B): the pairwise correlations (spearman correlation) between 59 exemplary traits in 734 segregants.

**Figure S4.**
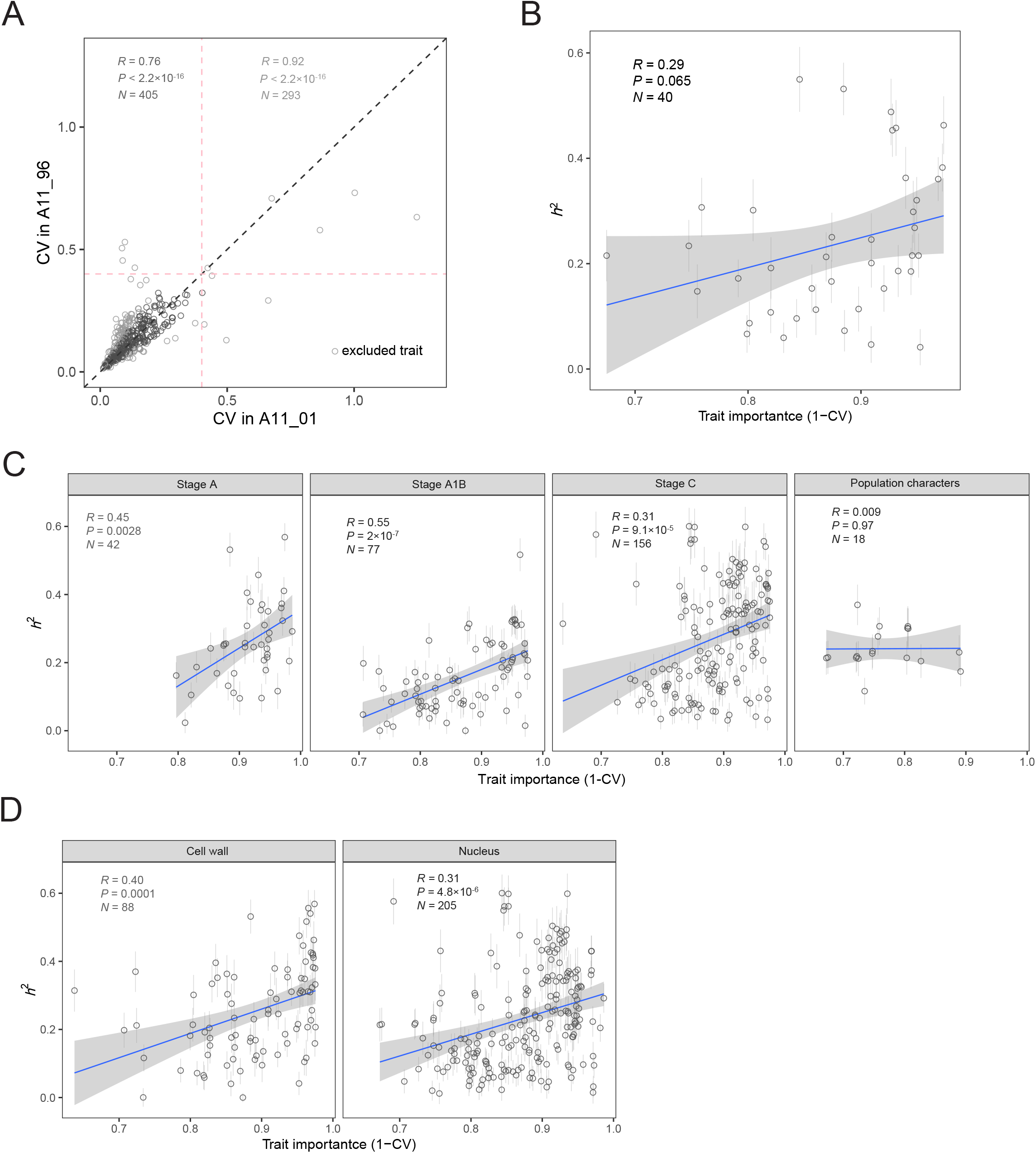
A positive relationship between trait importance estimated by 1-CV and *h*^2^. (A): the values of CVs of each morphological trait calculated by replications in A11_01 and A11_96. The light gray circles are excluded traits whose CV distances are larger than 0.2 or mean CVs are larger than 0.4. The gray dashed line is *y* = *x*, and the pink dashed lines represent *x* = 0.4 and *y* = 0.4, respectively. (B): the positive relationship between 1-CV and *h*^2^ among 59 exemplary traits. (C): the positive relationships between 1-CV and *h*^2^ were supported by traits in stage A, A1B, and C. (D): the positive relationships between 1-CV and *h*^2^ were supported by traits characterized different cell components.

**Figure S5.**
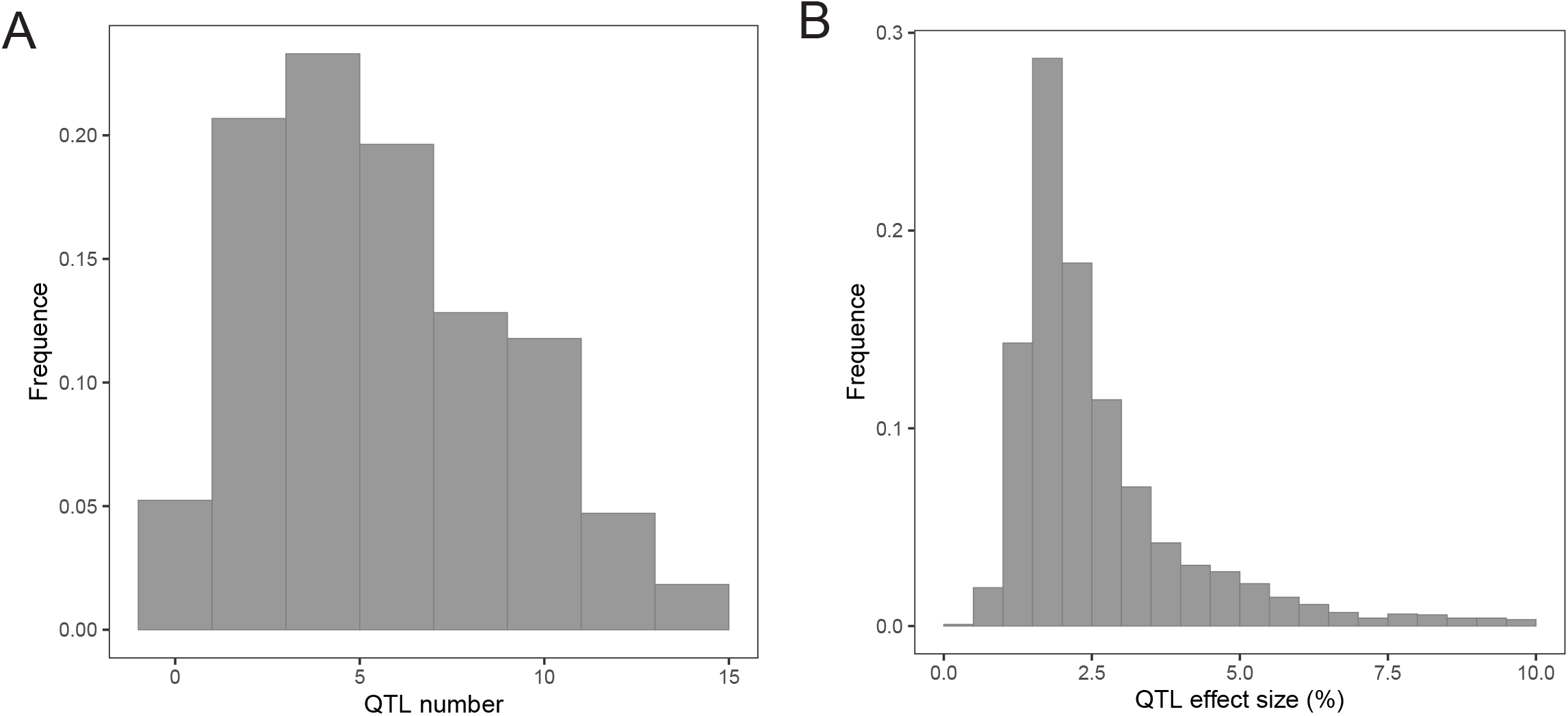
The distribution of QTL numbers detected for each morphological trait (A) and QTL effect size (B).

**Figure S6.**
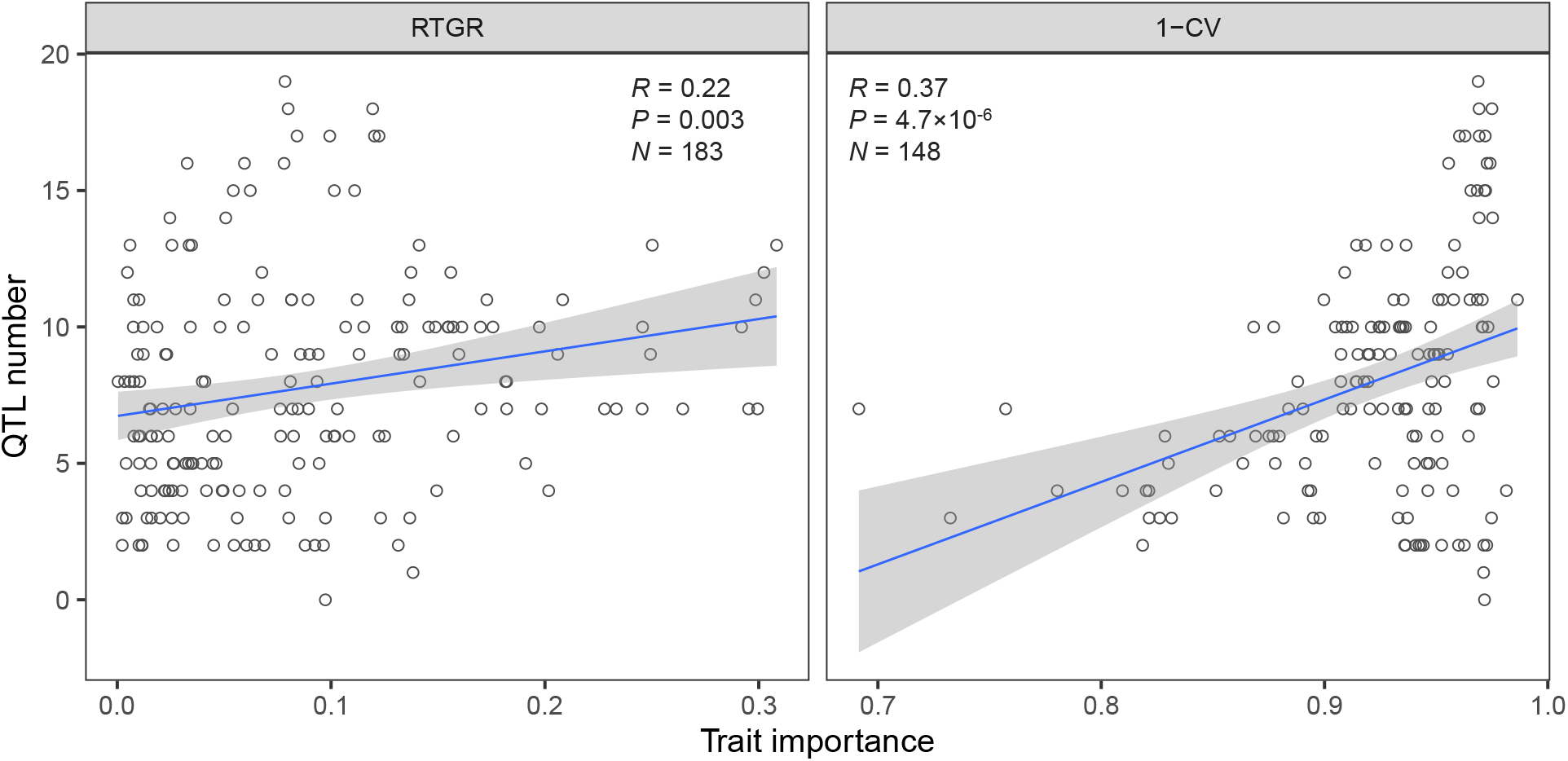
Positive correlations between QTL number and trait importance estimated by RTGR (left) and 1-CV (right) largely remain when only traits with available *f*_gene_ value were considered. The number of traits with *f*_gene_ value are 183 and 148 for RTGR and 1-CV, respectively.

## Legends for supplementary tables

**Table S1.** Raw values of morphological traits of 734 segregants measured in this study.

**Table S2.** Summary of the analyses of heritability and trait importance of 405 morphological traits in this study.

**Table S3.** Average value of normalized growth rate of each segregant.

**Table S4.** Summary of detected QTLs for each trait.

## References

Bates D, Machler M, Bolker BM, Walker SC. 2015. Fitting Linear Mixed-Effects Models Using lme4. Journal of Statistical Software 67:1–48.

Bloom JS, Ehrenreich IM, Loo WT, Lite TL, Kruglyak L. 2013. Finding the sources of missing heritability in a yeast cross. Nature 494:234–237.

Bloom JS, Kotenko I, Sadhu MJ, Treusch S, Albert FW, Kruglyak L. 2015. Genetic interactions contribute less than additive effects to quantitative trait variation in yeast. Nature Communications 6. preprocessCore: A collection of pre-processing functions [Internet]. https://github.com/bmbolstad/preprocessCore2019. Available from: https://github.com/bmbolstad/preprocessCore

Burger R, Gimelfarb A. 2002. Fluctuating environments and the role of mutation in maintaining quantitative genetic variation. Genet Res 80:31–46.

Chen H, Wu CI, He X. 2017. The genotype-phenotype relationships in the light of natural selection. Mol Biol Evol.

Crow JF. 2008. Maintaining evolvability. J Genet 87:349–353.

Crow JF. 2002. Perspective: Here’s to Fisher, additive genetic variance, and the fundamental theorem of natural selection. Evolution 56:1313–1316.

Endelman JB. 2011. Ridge Regression and Other Kernels for Genomic Selection with R Package rrBLUP. Plant Genome 4:250–255.

Frey BJ, Dueck D. 2007. Clustering by passing messages between data points. Science 315:972–976.

Fu Q, Posth C, Hajdinjak M, Petr M, Mallick S, Fernandes D, Furtwangler A, Haak W, Meyer M, Mittnik A, et al. 2016. The genetic history of Ice Age Europe. Nature 534:200–205.

Hellenthal G, Busby GBJ, Band G, Wilson JF, Capelli C, Falush D, Myers S. 2014. A genetic atlas of human admixture history. Science 343:747–751.

Hendry AP, Schoen DJ, Wolak ME, Reid JM. 2018. The Contemporary Evolution of Fitness. Annual Review of Ecology, Evolution, and Systematics, Vol 49 49:457–476.

Ho WC, Ohya Y, Zhang J. 2017. Testing the neutral hypothesis of phenotypic evolution. Proc Natl Acad Sci U S A 114:12219–12224.

Ho WC, Zhang J. 2014. The genotype-phenotype map of yeast complex traits: basic parameters and the role of natural selection. Mol Biol Evol 31:1568–1580.

Kosova G, Abney M, Ober C. 2010. Colloquium papers: Heritability of reproductive fitness traits in a human population. Proc Natl Acad Sci U S A 107 Suppl 1:1772–1778.

Kruuk LE, Clutton-Brock TH, Slate J, Pemberton JM, Brotherstone S, Guinness FE. 2000. Heritability of fitness in a wild mammal population. Proc Natl Acad Sci U S A 97:698–703.

Liedvogel M, Akesson S, Bensch S. 2011. The genetics of migration on the move. Trends Ecol Evol 26:561–569.

Merila J, Sheldon BC. 1999a. Genetic architecture of fitness and nonfitness traits: empirical patterns and development of ideas. Heredity 83:103–109.

Merila J, Sheldon BC. 1999b. Genetic architecture of fitness and nonfitness traits: empirical patterns and development of ideas. Heredity (Edinb) 83 (Pt 2):103–109.

Merila J, Sheldon BC. 2000. Lifetime Reproductive Success and Heritability in Nature. Am Nat 155:301–310.

Milot E, Mayer FM, Nussey DH, Boisvert M, Pelletier F, Reale D. 2011. Evidence for evolution in response to natural selection in a contemporary human population. Proc Natl Acad Sci U S A 108:17040–17045.

Mousseau TA, Roff DA. 1987. Natural selection and the heritability of fitness components. Heredity (Edinb) 59 (Pt 2):181–197.

Ohya Y, Sese J, Yukawa M, Sano F, Nakatani Y, Saito TL, Saka A, Fukuda T, Ishihara S, Oka S, et al. 2005. High-dimensional and large-scale phenotyping of yeast mutants. Proc Natl Acad Sci U S A 102:19015–19020.

Orr HA. 2009. Fitness and its role in evolutionary genetics. Nat Rev Genet 10:531–539.

Pettay JE, Kruuk LE, Jokela J, Lummaa V. 2005. Heritability and genetic constraints of life-history trait evolution in preindustrial humans. Proc Natl Acad Sci U S A 102:2838–2843.

Pizzo A, Roggero A, Palestrini C, Moczek AP, Rolandoa A. 2008. Rapid shape divergences between natural and introduced populations of a horned beetle partly mirror divergences between species. Evolution & Development 10:166–175.

Pulido F. 2007. The Genetics and Evolution of Avian Migration. BioScience 57:165–174.

Reale D, Festa-Bianchet M. 2000. Quantitative genetics of life-history traits in a long-lived wild mammal. Heredity (Edinb) 85:593–603.

Roy J, Arandjelovic M, Bradley BJ, Guschanski K, Stephens CR, Bucknell D, Cirhuza H, Kusamba C, Kyungu JC, Smith V, et al. 2014. Recent divergences and size decreases of eastern gorilla populations. Biology Letters 10.

Stirling DG, Reale D, Roff DA. 2002. Selection, structure and the heritability of behaviour. Journal of Evolutionary Biology 15:277–289.

Sved JA, McRae AF, Visscher PM. 2008. Divergence between Human Populations Estimated from Linkage Disequilibrium. American Journal of Human Genetics 83:737–743.

Sztepanacz JL, McGuigan K, Blows MW. 2017. Heritable Micro-environmental Variance Covaries with Fitness in an Outbred Population of Drosophila serrata. Genetics 206:2185–2198.

Teplitsky C, Mills JA, Yarrall JW, Merila J. 2009. Heritability of fitness components in a wild bird population. Evolution 63:716–726.

Wheelwright NT, Keller LF, Postma E. 2014. The effect of trait type and strength of selection on heritability and evolvability in an island bird population. Evolution 68:3325–3336.

Zhang XS. 2012. Fisher’s geometrical model of fitness landscape and variance in fitness within a changing environment. Evolution 66:2350–2368.

